# CTCF looping is established during gastrulation in medaka embryos

**DOI:** 10.1101/454082

**Authors:** Ryohei Nakamura, Yuichi Motai, Masahiko Kumagai, Haruyo Nishiyama, Neva C. Durand, Kaori Kondo, Takashi Kondo, Tatsuya Tsukahara, Atsuko Shimada, Erez Lieberman Aiden, Shinichi Morishita, Hiroyuki Takeda

**Author notes:** Equal contributions (Ryohei Nakamura, Yuichi Motai). Correspondence to; H.T., S.M., E.L.A.

## Abstract

Genome architecture plays a critical role in gene regulation, but how the structures seen in mature cells emerge during embryonic development remains poorly understood. Here, we study early development in medaka (the Japanese killifish, *Oryzias latipes*) at 12 time points before, during, and after gastrulation which is the most dramatic event in early embryogenesis, and characterize transcription, protein binding, and genome architecture. We find that gastrulation is most associated with drastic changes in genome architecture, including the formation of the first loops between sites bound by the insulator protein CTCF and great increase in the size of contact domains. However, the position of CTCF is fixed throughout medaka embryogenesis. Interestingly, genome-wide transcription precedes the emergence of mature domains and CTCF-CTCF loops.

## Main Text

Genome architecture plays a critical role in gene regulation, but how genome architecture is established during vertebrate development remains poorly understood. Embryogenesis itself consists of a highly orchestrated cascade of genetically encoded events, beginning with the fusion of the haploid male and female pronuclei. At first, the zygotic genome is transcriptionally silent, and rapid and synchronous cell cycles proceed under the control of maternally provided factors (*1*). During this process, epigenetic modifications and chromatin accessibility are globally reprogrammed (*2*). Next, zygotic genome activation (ZGA), simultaneous transcription of thousands of zygotic genes, occurs, and development proceeds under the control of zygotic gene products (*2*–*5*). Subsequently, gastrulation begins, as the cells in the single-layered blastula differentiate into three germ layers, and rudimentary organs begin to form (*6*). Thus, by the end of gastrulation, the vertebrate body plan is established.

Although extensive progress has been made by studying embryogenesis in mammals, fish, in particular medaka (the Japanese killifish, *Oryzias latipes*), have many advantages for embryological and genome studies, including high fecundity and *ex vivo* development (*7*). Medaka also benefit from a compact genome (roughly 800Mb) with an excellent assembly, comparable in quality with that of mouse (*8*). This greatly facilitates the generation of high-resolution contact maps, since the resolution varies inversely with the genome size. Here, we examined genome architecture, protein binding, and transcriptional activation before, during, and after gastrulation in medaka.

We began by examining the 3D genome structure of a medaka embryonic fibroblast cell line established from an embryo 4 days postfertilization (dpf) (referred to as mature fibroblasts) using *in situ* Hi-C (*9*), generating 952 million read pairs. The resulting matrices exhibited a typical plaid pattern associated with genome compartmentalization (Fig. 1A). Specifically, [i] all loci could be assigned to one of the two compartments, such that all loci in the same compartment exhibited an enhanced contact frequency with one another; [ii] contiguous chromatin intervals in the same compartment manifest as bright squares, called “contact domains,” along the diagonal of the contact map, i.e., domains in which all pairs of loci exhibit an enhanced contact frequency with one another; and [iii] comparison of the matrices with RNA-seq data confirmed that loci in one of the compartments are associated with active chromatin, whereas loci in the other compartment are associated with inactive chromatin (Fig. S1A). Such compartmentalization has been observed in many studies and species (*10*–*13*).

**Figure 1.**
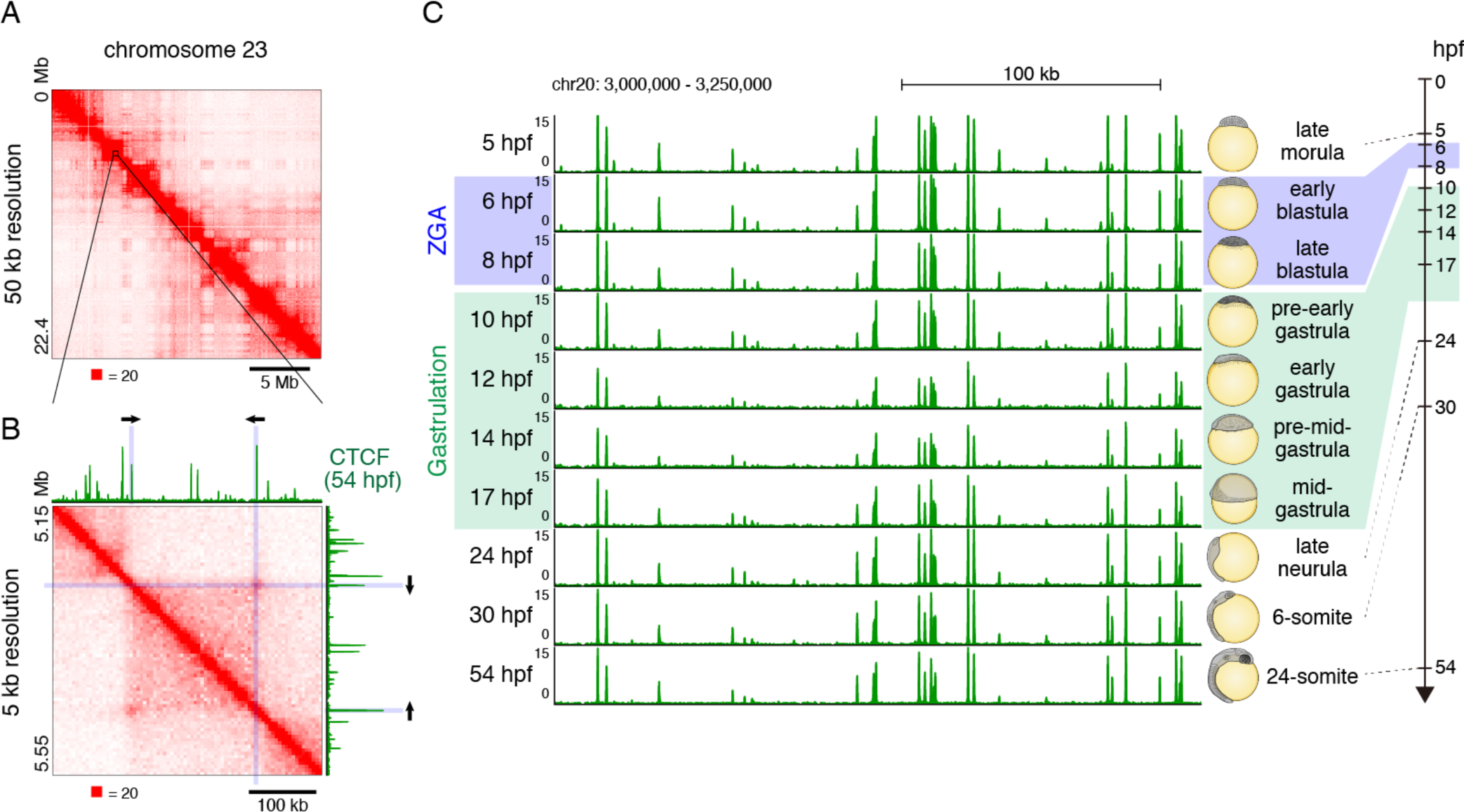
3D genome structure of medaka fibroblast cells and CTCF positioning throughout medaka development. (A) Observed Hi-C contact map of medaka fibroblast cells. Whole chromosome 23 at 50 kb resolution showing plaid pattern. (B) Zoomed view of chromosome 23 Hi-C contact map at 5 kb resolution and CTCF ChIP-seq track from 54 hpf embryos are shown. Arrows indicate the orientation of CTCF binding motif. (C) CTCF positioning is stable throughout medaka development; Representative view of CTCF ChIP-seq tracks across the developmental stages.

At higher resolutions, we observed numerous peaks in contact frequency indicating that a pair of anchor loci exhibit enhanced contact frequency with one another as compared to their neighbors along the contour of the chromosome (Fig. 1B, S1B). These peaks are consistent with the presence of point-to-point loops between pairs of anchor loci, and often demarcate contact domains. We also found that these peaks were closely associated with pairs of loci bound by the insulator protein CTCF, with the corresponding CTCF-bound motifs in the convergent orientation (i.e., pointing at one another) (Fig. 1B, S1C). The loop domains we observed in medaka closely resemble those seen in numerous studies in human and mouse. They are also consistent with a recent study in zebrafish, which found that CTCF motifs in the convergent orientation were enriched in the vicinity of contact domains (*10*). Taken together, these observations demonstrate that all of the features seen in mammalian Hi-C maps are also seen in medaka.

Next, we performed CTCF ChIP-seq experiments over a longitudinal time-course spanning 10 stages of medaka development: four time points before gastrulation, three during gastrulation, and three thereafter. Specifically, we sampled at stage 9 (late morula (256-512 cells), 5 hpf); stage 10 (early blastula (1k-cells), 6 hpf); stage 11 (late blastula (2-4k-cells), 8 hpf); stage 12 (pre-early gastrula, 10 hpf); stage 13 (early gastrula, 12 hpf); stage 14 (pre-mid-gastrula, 14 hpf); stage 15 (mid-gastrula, 17hpf); stage 18 (late neurula, 24 hpf); stage 21 (6-somite, 30 hpf); stage 27 (24-somite, 54 hpf). Crucially, this time-course is not possible in mammals, where the collection of a sufficient number of cells is prohibitive. No such data are available in zebrafish. We found that CTCF peak positions were extremely stable across all developmental stages examined (Fig. 1C, S2). Indeed, all CTCF motifs observed at loop anchor loci in mature fibroblasts were bound by CTCF as early as 5 hpf. This implies that, although CTCF may play a role in early genome architecture and regulation, changes in the positioning of CTCF are not a principal driver of the transcriptional and architectural changes seen during medaka embryonic development (see below).

We then examined the dynamics of genome architecture. For this, we performed in situ Hi-C (*9*) at twelve distinct stages. These include the 10 stages examined for CTCF mapping, as well as two additional intermediate stages (stage 10.5, 7 hpf; and stage 11.5, 9 hpf). The proportion of interphase cells were also assessed at early stages by microscope observation (Fig. S3). The Hi-C data was processed using Juicer (*14*) to obtain loop-resolution (5-10kb) *in situ* Hi-C maps for every time point studied (Fig. 2, S4). We found that neither compartments nor domains were present in the 5 hpf sample. Instead, the heatmap was a bright, undifferentiated diagonal, indicating the absence of any local features of genome architecture (Fig. 2). Starting at 7 hpf – the middle of zygotic genome activation – we observed the emergence of a plaid pattern, consistent with the presence of two compartments. This pattern became increasingly intense at each time point until 10 hpf (Fig. 2, S5). When we profiled open chromatin during the surrounding time points (5 hpf – 10 hpf) using ATAC-Seq, as well as ChIP-Seq using antibodies for H3K27 acetylation, we found that these features also appear at 7 hpf, and that their positioning was strongly associated with one of the two long-range patterns seen in the Hi-C map (Fig. S5). Overall, our findings are consistent with a recent study showing that the formation of open chromatin accompanies the emergence of compartmentalization and the activation of the zygotic genome in mice (*15*, *16*).

**Figure 2.**
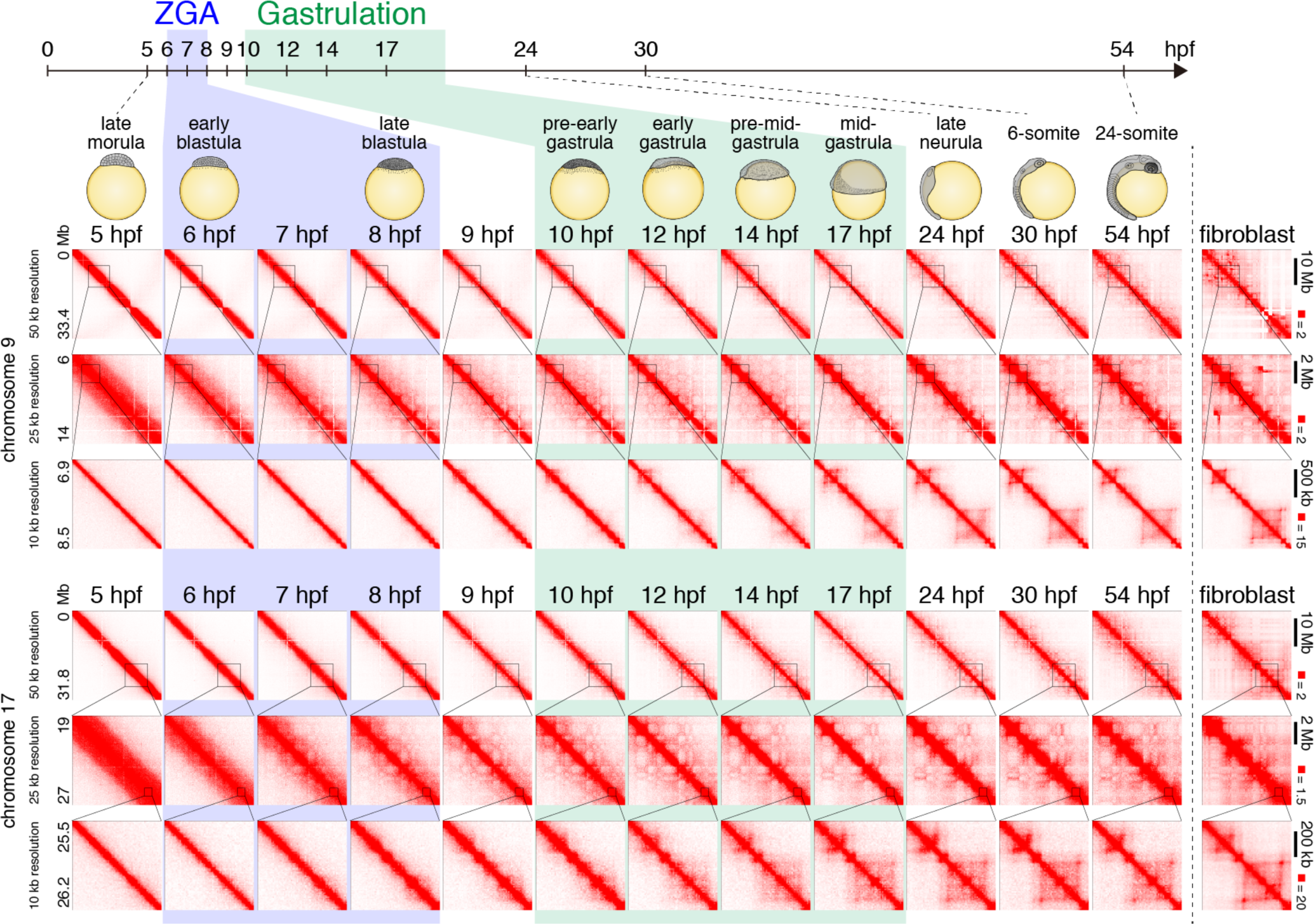
Chromatin contact dynamics across medaka development. Representative examples of Hi-C contact matrices from 12 time points across medaka development and fibroblast cells. Examples from chromosome 9 and 17 are shown for indicated resolutions. Note that chromosome 9 in fibroblast cells have chromosomal rearrangements.

Next, we looked for contact domains. We found that contact domains also began to be visible in our data at roughly 7 to 10 hpf (Fig. S6, S7). These domains were mostly small and relatively ambiguous as they were close to the diagonal. However, the size of domains increased greatly during gastrulation, and the domains at 24 hpf (late neurula) were much larger than those seen at 8 hpf (late blastula) (Fig. 3A, S10A). To quantify this effect, we calculated an N10 statistic for the domain size. This was calculated by examining all regions of the genome that are covered by a domain, and determining the minimum domain size required to cover 10% of these regions. (Thus, the remaining 90% are covered by domains that are smaller than the N10 value.) Thus, the N10 statistic reflects the size of the largest contact domains observed. The domain N10 at 7hpf, when we first observed domains, was 180 kb. This value did not change until the early gastrulation, and the domain N10 grew during gastrulation to 410 kb (24 hpf) (Fig. 3A). Notably, the N10 in mature fibroblasts was 465 kb, suggesting that the large contact domains associated with mature cells emerge during, but not before, gastrulation (Fig. S8).

**Figure 3.**
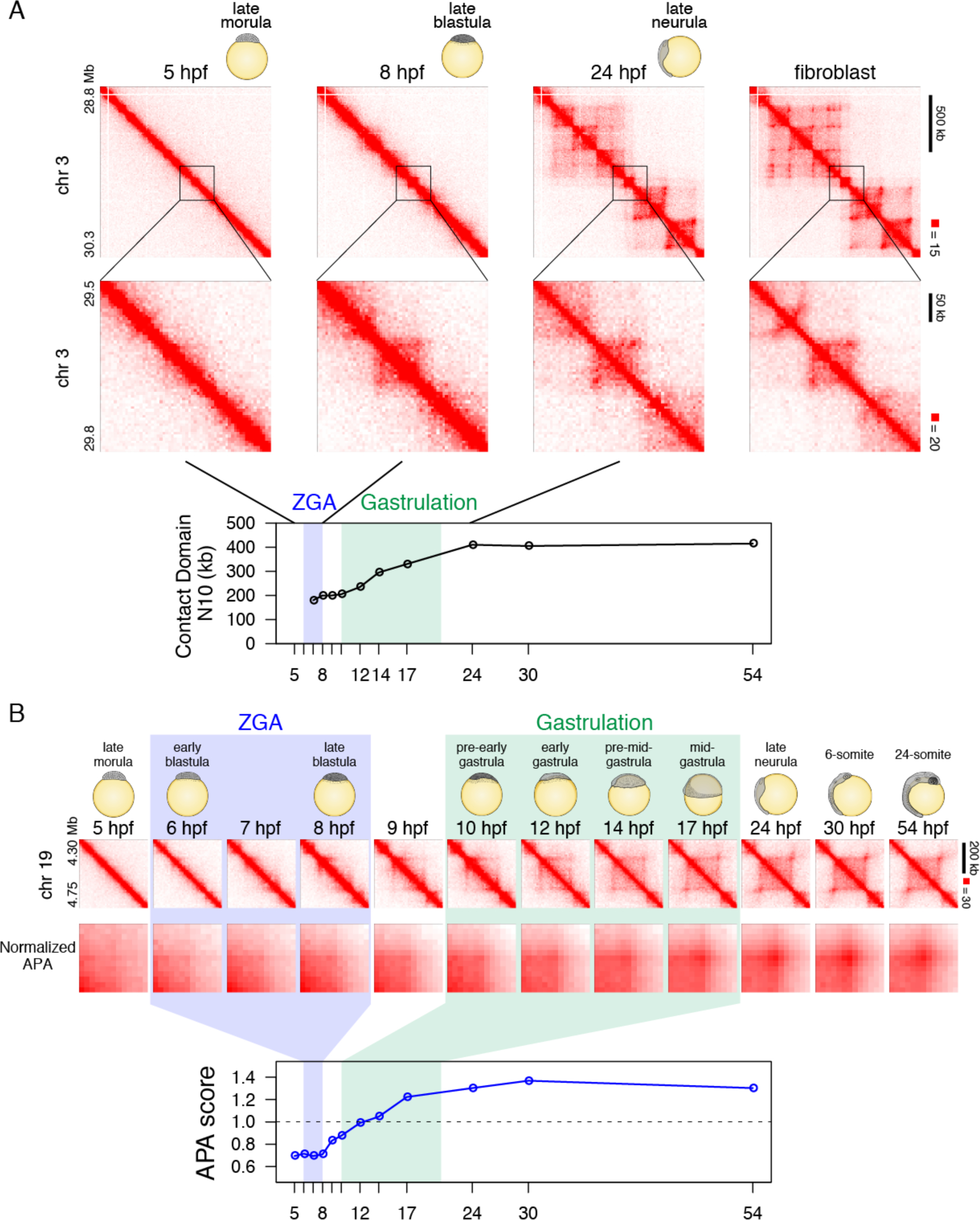
Establishment of CTCF-mediated loop domains during medaka development. (A) An example of establishment of large loop and small loop, and contact domain N10 statistics over stages are shown. Higher N10 values mean that larger contact domains cover longer intervals of genome. (B) Loops are established during gastrulation. Representative Hi-C contact map view of loop domain establishment is shown in upper portion. Normalized APA plots in middle portion show that loops become evident during gastrulation. APA scores (loop signal enrichment compared to lower-right signal of APA plot) across the developmental stages are shown in lower portion. Values greater than 1 indicate the presence of loops. Hi-C contacts around loops identified in fibroblast cells were aggregated for each stage.

Surprisingly, we did not observe the majority of loops in the data until gastrulation (Fig. 3A, 3B, S10A). Because automatic detection of loops suffered from many false positives due to higher noise near the diagonal, we filtered them out and counted the remaining reliable loops, i.e. loops associated with contact domains (*9*), and 100 kb or larger in size (See Supplemental Online Material). Thus, short-range (< 100 kb) loops, if any, were not subjected to further study. This method detected only two loops at 5 hpf (late morula) and 16 loops at 8 hpf (late blastula), whereas the number increased during gastrulation and to 405 at 24 hpf (Fig. S10B).

To rule out the possibility that loops were present at earlier time points, but were too weak to be detected individually, we performed Aggregate Peak Analysis (APA) (*14*) to examine whether the 1,869 loops (which are 100 kb or larger) observed in mature fibroblasts were reflected in maps from earlier timepoints. APA aggregates the Hi-C signal from a set of peaks: the center of the APA plot indicates the aggregate signal for the peak set, whereas adjacent values indicate the result of translating the peak set in various directions. Thus, an enhanced contact frequency in the central pixel of an APA plot indicates the presence of a signal from the loop list, in aggregate. Strikingly, we found no enrichment in the central pixel until 17 hpf (mid-gastrula) (Figure 3B). These results indicate that, despite consistent patterns of CTCF binding throughout development, point-to-point loops between CTCF bound sites do not form until 17 hpf.

Importantly, our findings imply that although zygotic genome activation and the onset of gastrulation may require the presence of CTCF, they do not require the formation of point-to-point loops between CTCF-bound sites. However, CTCF-mediated loops have been associated with gene transcription in numerous studies (*9*, *17*–*19*). To explore this relationship further, we performed RNA-seq for each of the 10 time points. Thus, we created a profile showing expression of every gene over time. To eliminate any confound from maternal transcripts that do not arise from zygotic transcription, we excluded all genes detectable at 5 hpf. We then compared the expression timing of genes that were associated with loops (i.e., the TSS was in the same 5kb bin as a CTCF loop anchor) in mature fibroblasts. We found that loop-associated genes tend to be expressed later in development than genes that are not associated with loops (Fig. 4A, 4B).

**Figure 4.**
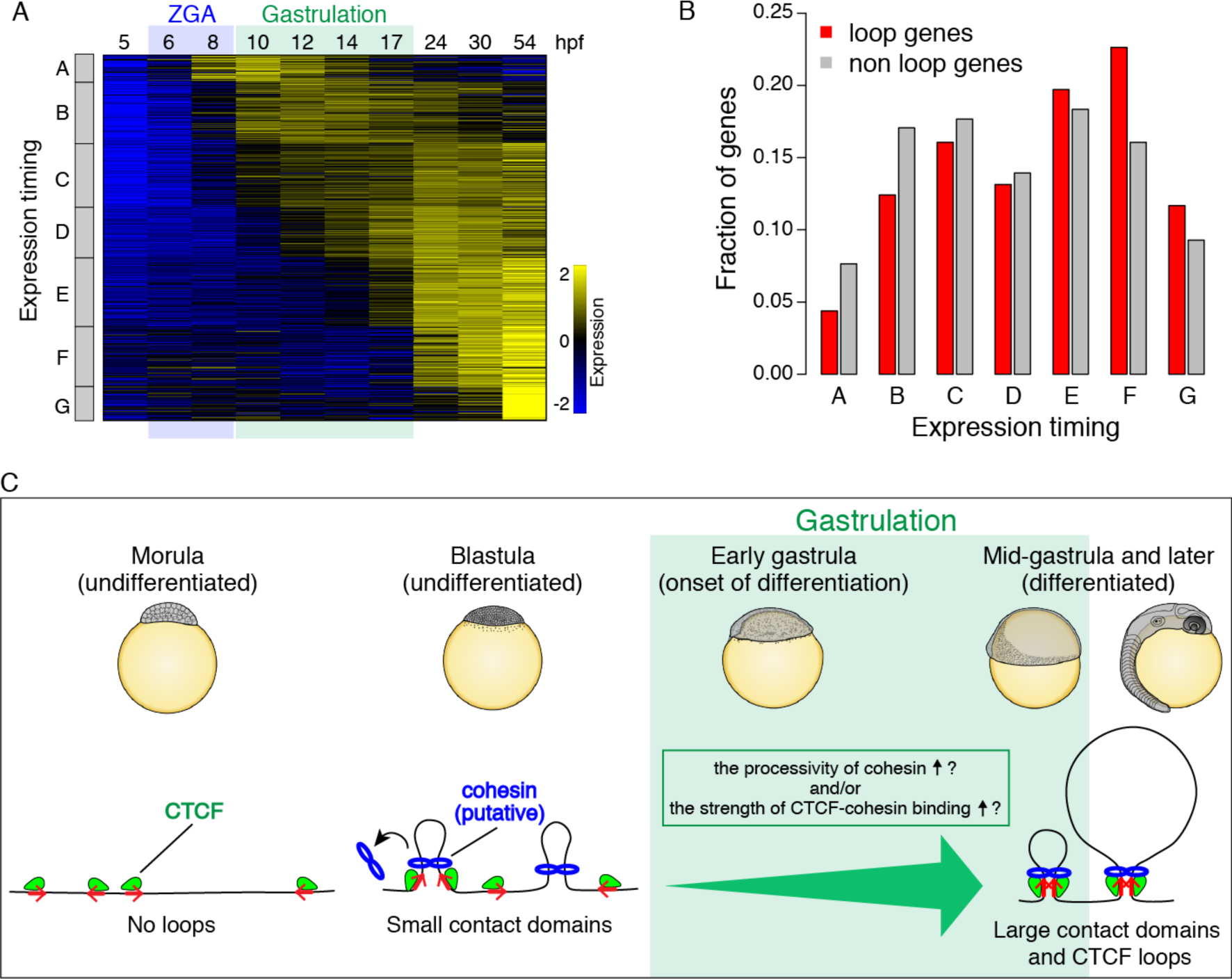
CTCF-mediated loops are associated with late activating genes. (A) Classification of zygotically expressed genes by expression dynamics. (B) loop genes are enriched with genes activated from mid-gastrula (17 hpf) or later (gene class E, F, and G). (C) A model of CTCF-mediated loop domains during medaka development.

This study provides a detailed overview of the establishment and maturation of genome architecture during embryogenesis, profiling the distribution of CTCF and the emergence of loops and domains before, during, and after gastrulation. We find that gastrulation is a critical event associated with numerous changes in genome architecture. Although we find that small contact domains emerge during ZGA (consistent with earlier studies), they become much larger during gastrulation, matching the sizes seen in mature cells. We also find that CTCF-CTCF loops first form during gastrulation, giving rise to loop domains.

Why do loops and domains suddenly form? The simplest model is that the distribution of CTCF changes, since its binding sites demarcate both loops and domains. Testing this hypothesis in mammals is challenging, because of the difficulty in routinely procuring adequate numbers of cells across a developmental timeline. By contrast, such experiments are straightforward in medaka. Surprisingly, we find that the distribution of CTCF is largely unchanged during differentiation. This implies that, at early stages in vertebrate development, CTCF is bound, and cohesin is present (Fig. S11), but loops do not form.

It is interesting to consider these findings in light of the loop extrusion model. In this model, loop domains form when a cohesin-based extrusion complex comprising two physically tethered subunits lands on chromatin, and the two subunits slide in opposite directions until they arrive at an inward-pointing CTCF site. The model suggests that, when cohesin is present, loops between CTCF sites should form spontaneously. This correspondence has been seen in all prior studies in which loop-resolution Hi-C and CTCF ChIP-seq data were generated. Thus, one possibility that accounts for the rapid and dramatic increase in domain size during gastrulation is a result of a sudden increase in the processivity of the loop extrusion complex. Similarly, the emergence of CTCF-CTCF loops during gastrulation might be driven by an increase in the strength of CTCF-cohesin binding (Fig. 4C). The factors that might be responsible for such changes will be an important topic for future studies.

Crucially, the late emergence of mature domain structures, including loop domains, suggests that they are not needed for initiating and regulating genome-wide transcription (i.e. ZGA). This finding is reminiscent of previous studies exploring the role of CTCF and cohesin disruption in mammalian cell lines, which have shown that the transcriptional consequences of such treatments are modest (*20*, *21*). The subsequent emergence of these features instead raises the possibility that loop domains play a role in the process of differentiation and lineage commitment.

## Acknowledgments

We thank H. Koseki for helpful discussion; M. Shamin for assistance with Juicer parameter setting; and Y. Yamagishi for assistance with animal husbandry and embryo sampling.

## Author contributions

R.N. conceived the project. Y.M. found loops in fibroblast to explore analysis in gastrulation. R.N. and Y.M. analyzed data. R.N. performed embryonic Hi-C and RNA-seq experiments. M.K. performed fibroblast Hi-C and RNA-seq experiments. R.N. and H.N. performed ChIP-seq experiments. T.K., K.K. and A.S. performed 3D-FISH experiments. T.T. designed CTCF antibody. M.K. analyzed d-rR/HNI hybrid RNA-seq data. N.C.D. assisted in analyzing Hi-C data. H.T., S.M., and E.L.A supervised the project. R.N., Y.M., H.T., S.M., and E.L.A. prepared the manuscript with input from all authors.

## Data and materials availability

All sequencing data are available in the DDBJ Sequence Read Archive (DRA) upon publication.

## Materials and Methods

### Fish strains and sampling

We used medaka d-rR wild-type strain. For embryonic RNA-seq data, we used female HNI-II strain and male d-rR strain, and obtained F1 hybrid medaka embryos. Medaka fishes were maintained and raised under standard condition. Embryos were dechorionated and incubated at 28°C. Developmental stages were determined based on a previous study (*22*). Medaka embryonic fibroblast cell line was previously established in our laboratory. All experimental procedures and animal care were carried out according to the animal ethics committee of the University of Tokyo (Approval No. 14-05).

### *in situ* Hi-C

*in situ* Hi-C experiments were performed as previously described (*9*) with some modifications. For all developmental stages, two biological replicates were generated. Three biological replicates were generated for fibroblast cells. Dechorionated embryos were collected in L-15 medium with 15% FBS, and dissociated by pushing through a 21G needle using a syringe. Dissociated embryos or fibroblast cells were collected by centrifuge at 500 x g for 3 minutes, and cross-linked with 1% formaldehyde in 1.5 ml L-15 medium (15% FBS) for 10 minutes at room temperature. Cross-linking was quenched by adding 2.5M glycine (125 mM final) and incubating for 5 minutes at room temperature, then 15 minutes on ice. Cells were pelleted at 500 x g for 5 minutes, supernatant was removed, and stored at −80°C.

Cells were thawed on ice, washed with PBS, resuspended in 250 µl of ice-cold Hi-C lysis buffer (10 mM Tris-HCl pH8.0, 10 mM NaCl, 0.2% Igepal CA-630) with 50 µl of protease inhibitors (Sigma, P8340), and incubated on ice for 20 minutes. The lysate was centrifuged at 2,500 x g for 5 minutes, washed with ice-cold Hi-C lysis buffer, resuspended in 50 µl of 0.5% SDS, and incubated for 10 minutes at 62°C. 145 µl of water and 25 µl of 10% Triton X-100 were added and incubated for 15 minutes at 37°C. 25 µl of NEBuffer2 and 100 U of MboI were added, and chromatin was digested for overnight at 37°C with rotation. MboI was inactivated by incubating at 62°C for 20 minutes. DNA ends were labeled with biotin by adding 50 µl of fill-in master mix (37.5 µl of 0.4 mM biotin-14-dATP (Life Technologies), 1.5 µl of 10 mM dCTP, 1.5 µl of 10 mM dGTP, 1.5 µl of 10 mM dTTP, 8 µl of 5 U/µl DNA Polymerase I Large Klenow Fragment (NEB)), and incubated at 37°C for 1.5 hours with rotation. Proximal ligation was performed by adding 900 µl of ligation master mix (669 µl of water, 120 µl of 10x NEB T4 DNA ligase buffer, 100 µl of 10% Triton X-100, 6 µl of 10 mg/ml BSA, and 5 µl of 400 U/µl T4 DNA ligase (NEB)), and incubated at room temperature for 4 hours. Proteins were degraded by adding 50 µl of 20 mg/ml proteinase K, 120 µl of 10% SDS, and incubated at 55°C for 2 hours. Cross-linking was reversed by adding 130 µl of 5M NaCl and incubated at 68°C for overnight.

The biotinylated DNA was collected by ethanol precipitation, and resuspended in 130 µl of Tris buffer (10 mM Tris-HCl, pH 8.0). DNA was sheared using Covaris S220 with following parameters; Peak Incident Power: 140, Duty Factor: 10, Cycle per Burst: 200, time: 80 seconds. The sheared DNA was size selected to a size of 100-500 bp and purified using AMPure XP beads (Bechman Coulter). DNA was eluted in 60 µl of Tris buffer.

The sheared DNA was end-repaired by adding 40 µl of master mix (10 µl of 10x NEB T4 DNA ligase buffer with 10 mM dATP, 20 µl of 2.5 mM dNTP, 5 µl of 10 U/µl NEB T4 PNK, 4 µl of 3 U/µl NEB T4 DNA polymerase I, 1 µl of NEB DNA polymerase I Large Klenow Fragment), and incubated at room temperature for 30 minutes. End-repaired DNA was then purified using MinElute PCR purification Kit (Qiagen) and eluted in 40 µl of water. DNA was A-tailed by adding 20 µl of master mix (10 µl of 10x NEBuffer 2, 5 µl of 10 mM dATP, 5 µl of 5 U/µl NEB Klenow exo minus), and incubated at 37°C for 30 minutes. The sample was then incubated at 65°C for 20 minutes, and 200 µl of water was added.

150 µl of 10 mg/ml Dynabeads MyOne Streptavidine T1 beads (Life technologies) was washed with 400 µl of 1x Tween Washing Buffer (1x TWB: 5 mM Tris-HCl pH7.5, 0.5 mM EDTA, 1 M NaCl, 0.05% Tween 20), and resuspended in 300 µl of 2x Binding Buffer (2x BB: 10 mM Tris-HCl pH7.5, 1 mM EDTA, 2 M NaCl). The Beads were added to the A-tailed DNA sample, and incubated at room temperature for 15 minutes with rotation. The biotinated DNA-bound beads were collected with a magnet and supernatant was discarded. The beads were washed twice by adding 600 µl of 1x TWB, transferred to a new tube, incubated at 55°C for 2 minutes on a Thermomixer, the supernatant was discarded using a magnet. Then, the beads were washed with 100 µl of 1x Quick ligase buffer (NEB), resuspended in 50 µl of 1x Quick ligation buffer. 2µl of NEB DNA Quick ligase and 1.5 µl of 30 uM indexed Illuminar adapter were added, and incubated at room temperature for 15 minutes. The beads were washed twice by adding 600 µl of 1x TWB, transferred to a new tube, incubated at 55°C for 2 minutes on a Thermomixer, the supernatant was discarded using a magnet. The beads were washed with 100 µl of Tris buffer, transferred to a new tube, and resuspended in 50 µl of Tris buffer. The libraries were amplified directly off of the beads with 6-7 cycles of PCR, using KAPA HiFi HotStart ReadyMix (KAPA Biosystems), and DNA was purified using AMPure XP beads. Paired-end sequencing of the Hi-C libraries were performed using the Illumina HiSeq 1500 platform.

### Hi-C data processing

The sequenced reads were mapped, filtered, and normalized using Juicer (version 1.5) (*14*). In order to compare between different stages, KR-normalized matrix was further normalized by average total contacts of each row, so that all samples will have a same total contact number. We used Juicer Tools (version 1.7.6) to calculate Pearson’s correlation matrices, eigenvectors, and others in what follows.

Loops were identified using Juicer Tools HiCCUPS at 5 and 10 kb resolution. As our fibroblast cells were maintained in a laboratory environment for a long period of time, chromosomes could have been rearranged, thereby yielding a number of peaks identified by HiCCUPS in the off-diagonal areas of the Hi-C contact maps. In particular, off-diagonal loops of >1Mb in size were unexpectedly long, were likely to be false-positive, and were therefore removed. We also noticed that skewed sequence coverage bins sometime caused abnormal enrichment on contact maps. To filter out these possibly false-positives loops, we selected loops on rows and columns of each contact matrix where the value of KR normalization vector produced for matrix balancing was in the range of 0.5 to 2.0.

As we did not have CTCF ChIP-seq data for fibroblast cells, as substitutes, we instead used CTCF ChIP-seq peaks identified from 54 hpf embryos. The CTCF motif orientation of loops were determined using MotifFinder of Juicer Tools. Specifically, we used motif MA0139.1_CTCF from Jasparcore 2014 as CTCF motif, and the positions of the CTCF motif in medaka genome were identified using FIMO (*23*) from MEME Suite with the default parameters.

Aggregate peak analysis (APA) was performed using Juicer Tools using all filtered fibroblast loops that are 100 kb or larger at 5 kb resolution by using command with: apa ‐n 20 ‐w 6 ‐q 4 ‐r 5000. In the presence of strong background signals near diagonal, we considered loops of size 100kb or larger (see the rationale in Fig. S9). Normalized APA plots were used to depict loop signal enrichment across stages. P2LL score (the ratio of the central pixel to the mean of the pixels in the lower left corner) from Juicer output was used for the quantification of loop strength.

Contact domains were annotated using the Juicer Tools Arrowhead algorithm at 5kB resolution, changing the default filter size from 300kB to 50kB. To quantify the change in size of contact domains during development, we used the N10 statistics (Fig. 3E), where the N10 maximizes such that non-overlapping contact domains of size N10 or longer occupy at least 10 percent of the genome covered by domains. To calculate the N10 value, one can order non-overlapping contact domains in the descending order and scan the ordered list from the top until the sum of domain sizes becomes 10% or larger. A higher N10 value means that larger contact domains occupy at least 10% of the genome covered by contact domains. The statistics show that contact domains with larger size are established later during gastrulation (Fig 3E).

### ATAC-seq

ATAC-seq was performed as previously described (*24*) with some modifications. Embryos were homogenized in PBS, and cells were harvested by centrifugation at 500 x g for 5 minutes. Approximately 5,000 cells were used for each experiment. After washing with PBS, cells were resuspended in 500 µl of cold lysis buffer (10 mM Tris-HCl pH 7.4, 10 mM NaCl, 3 mM MgCl_2_, 0.1% Igepal CA-630), centrifuged for 10 minutes at 500 x g, and supernatant was removed. Tagmentation reaction was performed as described previously (*24*) with Nextera Sample Preparation Kit (Illumina). After tagmented DNA was purified using MinElute kit (Qiagen), two sequential PCR were performed to enrich small DNA fragments. First, 9-cycle PCR were performed using indexed primers from Nextera Index Kit (Illumina) and KAPA HiFi HotStart ReadyMix (KAPA Biosystems), and amplified DNA was size selected to a size of less than 500 bp using AMPure XP beads (Bechman Coulter). Then a second 7-cycle PCR was performed using the same primer as the first PCR, and purified by AMPure XP beads. Libraries were sequenced using the Illumina HiSeq 1500 platform.

### RNA-seq

Embryos were homogenized in 1 ml of Isogen (Nippongene), and 200 µl of chloroform was added, and total RNA was isolated using RNeasy MinElute Cleanup Kit (Qiagen). Total RNA from Fibroblast cells was extracted using RNeasy Mini kit (Qiagen). Ribosomal RNA was removed using RiboMinus Eukaryote System v2 (Thermo Fisher Scientific), and RNA-seq libraries were generated using KAPA Stranded RNA-seq Kit (KAPA Biosystems). Libraries were sequenced using the Illumina HiSeq 1500 platform.

### ChIP-seq

ChIP was performed as previously described with modifications (*25*). Dechorionated embryos were dissociated using a 21G needle in PBS containing 20 mM Na-butyrate, complete protease inhibitor (Roche) and 1 mM PMSF, and fixed with 1% formaldehyde for 8 minutes at room temperature then quenched by adding glycine (200 mM final) and incubating on ice for 5 minutes. After washing with PBS, cell pellets were stored at −80°C. Cells were thawed on ice, suspended in lysis buffer (50 mM Tris-HCl (pH 8.0), 10 mM EDTA, 1% SDS, 20 mM Na-butyrate, complete protease inhibitors, 1 mM PMSF), sonicated ten times using sonifier (Branson) at power seven, and centrifuged for collecting chromatin lysates. The chromatin lysates were diluted with RIPA ChIP buffer (10 mM Tris-HCl (pH 8.0), 140 mM NaCl,1 mM EDTA, 0.5 mM EGTA, 1% Triton-X100, 0.1% SDS, 0.2% sodium deoxycholate, 20 mM Na-butyrate, complete protease inhibitors, 1 mM PMSF) and rotated with an antibody/protein A Dynabeads complex for overnight at 4°C. Immunoprecipitated samples were washed three times with RIPA buffer (10 mM Tris-HCl (pH 8.0), 140 mM NaCl,1 mM EDTA, 0.5 mM EGTA, 1% Triton-X100, 0.1% SDS, 0.2% sodium deoxycholate) and once with TE buffer, followed by elution with Lysis buffer at 65°C for overnight. Eluted samples were treated with RNase A for 2 hours at 37°C, proteinase K for 2 hours at 55°C, and DNA was purified by phenol/chloroform extraction and ethanol precipitation. Input DNA was simultaneously treated from the elution step. ChIP-seq libraries were generated using KAPA Hyper Prep Kit (KAPA Biosystems). Antibodies used in this study are as follows; H3K27ac: ab4729 (abcam) CTCF: see below.

### Generation of Medaka CTCF antibody

Four peptides derived from medaka CTCF (gene ID, ENSORLG00000008771) were synthesized by MAB Instirute, Inc (Nagano, Japan) as listed below.

~~~
1-36+C   METGQATALASDGKVLSEGGEALIQTGQGDEAGTMEC
~~~

~~~
50-70+C  MKTEVLEGGGTVTVTGGDEGQC
~~~

~~~
699-730  CTDETTEQVITGGGKPGAQSEELSQADAAAQE
~~~

~~~
C+734-753 CSAAPSNGDLTPEMILSMMDR
~~~

The mixture of these peptides was used as an antigen to generate antibodies against medaka CTCF in mice and hybridoma clones were established by MAB Institute, Inc. We screened hybridoma clones by western blotting using the lysate of HEK293 cells transfected with pCS2-MT-olCtcf1. For the selected clones, the supernatant of culture medium was concentrated by ammonium sulfate precipitation and antibodies were purified by protein A sepharose and eluted with 100mM Glycine, pH 4.0, then dialyzed with PBS + 0.05 % NaN_3_.

### ATAC-seq and ChIP-seq data processing

The sequenced reads were preprocessed to remove low-quality bases and adapter derived sequences using Trimmomatic v0.32 (*26*), and then aligned to the medaka reference genome version 2.2.4 by BWA (*27*). Reads with mapping quality (MAPQ) larger than or equal to 20 were used for the further analyses. For ATAC-seq and ChIP-seq, MACS2 (version 2.1.1.20160309) (*28*) was used to call peaks and generate signals per million reads tracks using following commands; ATAC-seq: macs2 callpeak ‐‐ nomodel ‐‐extsize 200 ‐‐shift ‐100 ‐g 600000000 ‐q 0.01 ‐B ‐SPMR, H3K27ac and CTCF ChIP-seq: macs2 callpeak ‐q 0.01 ‐B ‐SPMR. The option ‐g 600000000 is the effective genome size estimated by re-mapping 50 mer of medaka reference genome version 2.2.4 to itself using GEM library tools; gem-indexer (build 1.423), gem-mappability (build 1.315), and counting the uniquely remapped reads denoted as “!” in the output (*29*). The number of peaks called by MACS2 depends on total number of reads. Thus, to balance the total read number, 10 million reads were randomly sampled from each of the replicate for CTCF ChIP-seq data. For figure 1C and S2, replicates were combined for each stage after random sampling.

### RNA-seq data processing for d-rR embryos (5 to 54 hpf) and fibroblast cells

The sequenced reads were preprocessed to remove low-quality bases and adapter derived sequences using Trimmomatic v0.32 (*26*), and were aligned to the medaka reference genome version 2.2.4 by STAR (*30*) and reads with mapping quality (MAPQ) larger than or equal 20 were used for the further analyses.

### Analysis of RNA-seq data from d-rR/HNI hybrid F1 embryos (5 to 10 hpf)

To investigate the transcriptional activity in early zygotic stages, we employed hybrid F1 of d-rR and HNI strains, the former was used as a paternal strain and the later as a maternal strain, for RNA-seq analysis. We mapped RNA-seq reads of hybrid F1 embryos to both Hd-rR and HNI reference genomes (v2.2.4) (*31*). For reliable identification of the source strain of expressed RNA, paternal Hd-rR or maternal HNI, we utilized the genomic polymorphism between the two strains because the genetic divergence is sufficiently high and the single nucleotide variant (SNV) rate is ~2.5% (*31*). First, to call SNVs between the two genomes, we obtained publicly available whole genome shotgun sequencing data of HNI strain with accession numbers DRR002216 and DRR002217 from DDBJ. Raw Illumina reads were preprocessed to remove low-quality bases and adapter derived sequences using Trimmomatic v0.32 (*26*). Subsequently, a total of 109-fold coverage short read data were aligned with medaka Hd-rR reference genome (v2.2.4) by using BWA v0.7.8 (*27*). After refining alignments around indel sites with GATK v3.6 (*32*), SNVs and indels were called by samtools v1.2 (*33*). High quality variants, SNVs and indels, with read depth >= 20 and variant quality >= 20 were used for downstream analysis. To further filter erroneous regions where short reads from HNI matched to the Hd-rR genome, we aligned HNI blastula RNA-seq reads (SRR3168578) with the two genomes using STAR v2.5.4 (*30*), and discarded variant sites where HNI RNA-seq reads matched erroneously, retaining 17,516,984 sites for hybrid F1 embryo RNA-seq analyses. After mapping RNA-seq reads of hybrid F1 of each developmental stage the two genomes using STAR, we determined that reads were paternally expressed if they matched the Hd-rR sides of polymorphic SNVs. For Supplementary Figure S4A, paternal read coverage was normalized by the sum of paternal and maternal read numbers to calculate rpm.

### Alignment of compartments and profiles of ATAC-seq, RNA-seq and ChIP-seq

To compare compartments represented by two-dimensional patterns with one-dimensional profiles such as ATAC-seq, RNA-seq and ChIP-seq data, we used the one-dimensional eigenvectors of the Pearson’s correlation matrices of the Hi-C contact maps in 50 kb resolution (*34*). In earlier developmental stages, the compartmentalization occurred proximal to the diagonals of the contact maps (Fig. S5), and the global chromosome-wide eigenvector did not appear to represent individual local plaid patterns of compartmentalization. However, enlarging the chromosome-wide eigenvector, we found a strong correlation of respective positive and negative values in the sub-eigenvector with active and inactive states in compartmentalization. As an eigenvector is invariant up to a scalar multiplication, we negated a sub-eigenvector by multiplying it by −1 when the vector and the ATAC-seq profiles should be correlated positively.

### Gene expression at loop anchors

The RPKM values were computed from RNA-seq read counts for medaka gene model version 2.2.4 (http://utgenome.org/medaka_v2/#!Medaka_gene_model.md) in each time point. Out of medaka gene model v2.2.4, genes associated with Ensembl protein ID were used. Then, we defined zygotically expressed genes (ZEGs) such that the read counts < 30 and the RPKM < 2 at 5 hpf, and read counts > 30 and RPKM > 4 in one of later stages than 5 hpf, which means that ZEGs were inactive before ZGA and began to be transcribed during or after that. Then we got the vectors whose elements were the gene expression levels of ZEGs in developmental time course. To separate ZEGs by expression timing, the vectors were grouped into seven clusters according to a widely-used method (*35*). Loop genes in ZEGs were defined so that their TSSs and loop anchors with loop size > 100 kb (see the rationale in Fig. S9) were within the same 5 kb bin. Non-loop genes were the rest ZEGs.

### Association of large contact domains with mature cells

We compared contact domains between mature fibroblast cells and each time point during development. Two contact domains were treated identical when their boundaries were close, which was however a difficult task to do because the boundaries changed between samples. Indeed, the boundaries of 54 hpf and fibroblast (both were most matured samples in our data) should be consistent; however, only 8 % of boundaries at 54 hpf exactly matched those at fibroblast. To increase the ratio, we allowed the positional difference between pairs of matching domain boundaries to be at most 25 kb (5 bins) so that 71% domains at 54 hpf were associated with those at fibroblast. We counted the number of large contact domains which were shared with mature fibroblast cells, and plotted for three different domain size ranges; ≥ 200 kb, ≥ 300 kb, and ≥ 400 kb (Fig. S8).

### Detecting more reliable loops

Automatic detection of loops from Hi-C contact maps generally suffered from many false positives. To filter out false positive loops, we imposed two additional constraints on loops; namely, loops were associated with contact domains (Fig. S10), and loops were 100 kb or larger in size by considering the resolution limitation of identifying loops (Fig. S9). The first condition was supported by the fact that a large fraction of peaks in mammals were associated with domains (*9*), which was also the case in the medaka genome. To meet the first condition, we extracted loops (called by HICCUPS) associated with contact domains by selecting those located in boundaries of contact domains within a distance of 20 kb (4 bins).

### 3D-FISH

3D-FISH analyses were carried out following protocols described previously with some modifications (*36*). Medaka fish embryos were fixed with 4% paraformaldehyde (PFA), and embedded in paraffin and sectioned into 5 µm thickness. After deparaffinization, HistoVT ONE (Nakalai Tesque, 06380-05) treatment and wash process by water, hybridizations with Medaka BAC as probes were proceeded. BACs, ola1-199A02, ola1-190N08 and ola1-182E06 were labeled with ChromaTide^TM^ AlexaFluor^TM^ 488-5-dUTP (Thermo Fisher Scientific C11397), Cy3-dCTP (GE Healthcare PA53021) or Cy5-dCTP (GE Hearlthcare PA55021) by nick-translation (Roche 976776), respectively.

The confocal images were captured by microscopy Olympus UPlanSApo 100x NA 1.40 and PlanApo N 60x NA 1.42. Images with 65 nm pixels in X-Y and 300 nm steps in Z axes. Those images were deconvoluted and distances of foci (centroid) were measured by using ImageJ.

### Counting mitotic cell ratio in early embryos

Embryos were fixed with 4% PFA at room temperature for 2 hours, then 4°C for overnight. The fixed embryos were stained with DAPI, and mitotic cells were identified manually based on the morphology of condensed chromosomes. Cell number was counted using ImageJ.

**Fig. S1.**
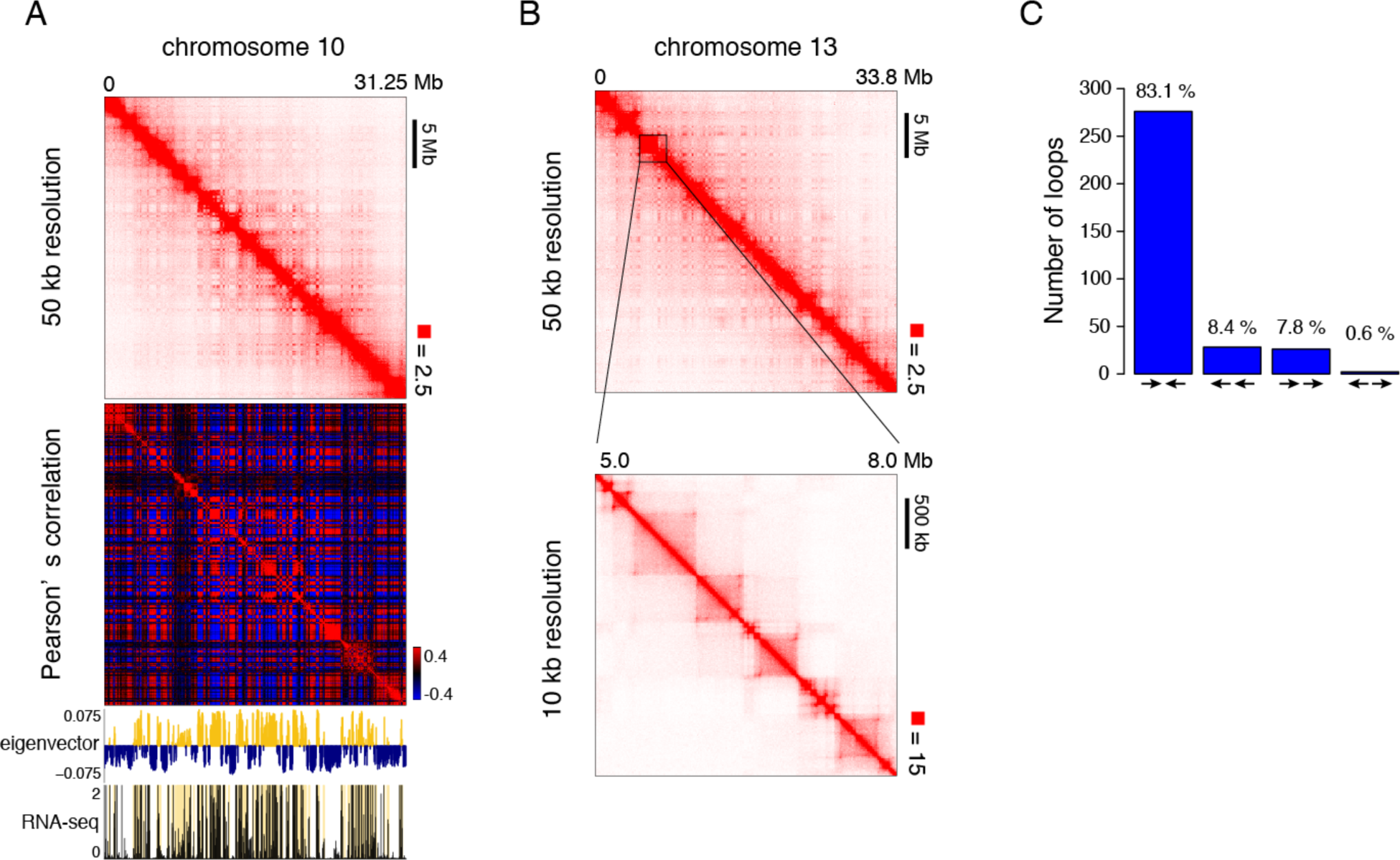
Compartments and CTCF loops organize 3D genome structure of medaka fibroblast cells. (A) From top to bottom, observed contact matrix of whole chromosome 10 at 50 kb resolution, Pearson’ s correlation matrix, eigenvector, and RNA-seq tracks are shown. The A compartments are highlighted with yellow in eigenvector and RNA-seq (rpkm) tracks. (B) An example of compartments and loops. Observed Hi-C contact matrix of medaka fibroblast cells. Whole chromosome 13 at 50 kb resolution showing plaid pattern (top) and zoomed view at 10 kb resolution showing loop domains. (C) Number of loops with indicated combination of unique CTCF motif orientation.

**Fig. S2.**
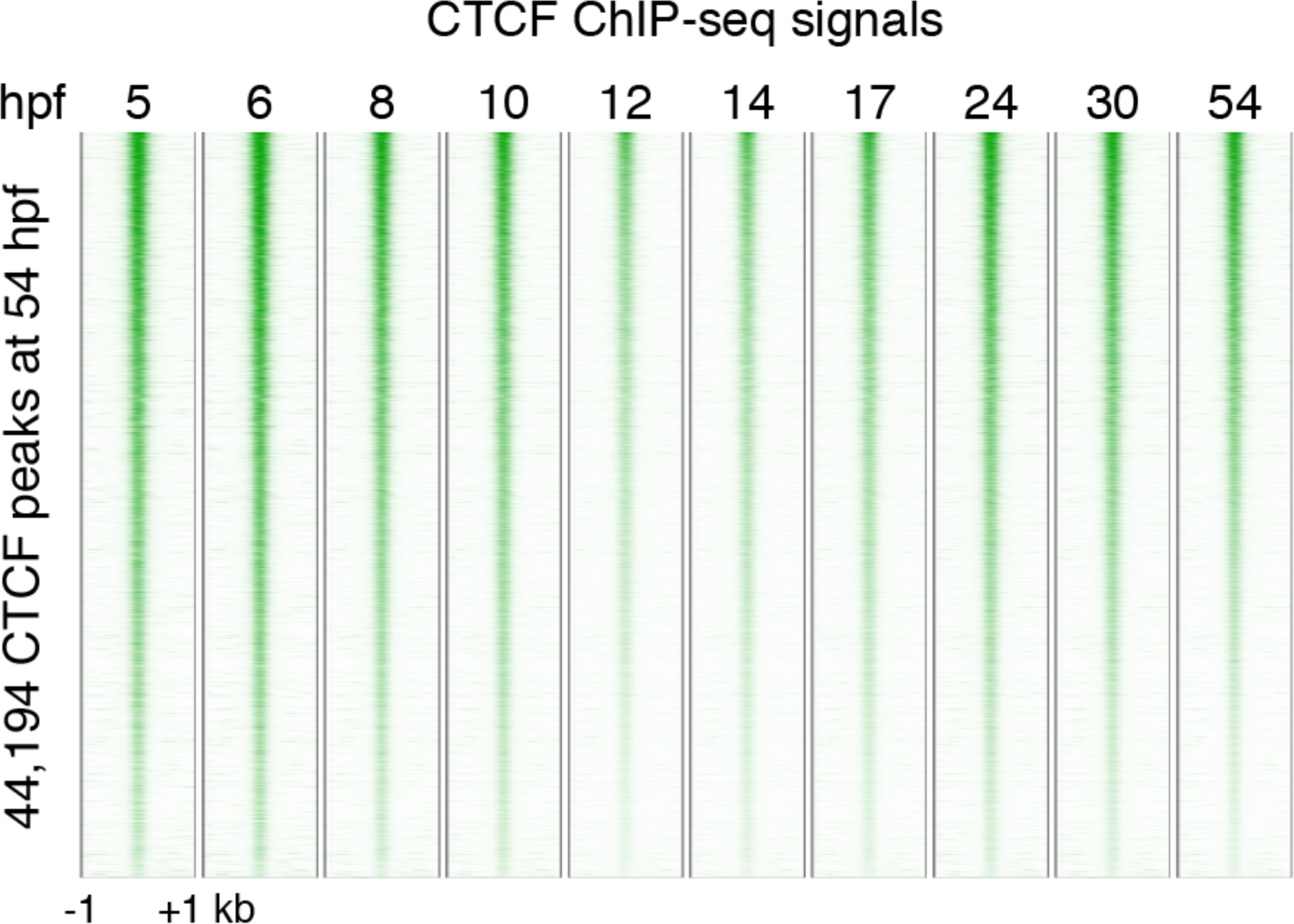
CTCF binds stably across all developmental stages. CTCF ChIP-seq signals aligned to the positions of the peaks at 54 hpf. Rows correspond the 44,194 CTCF signal peaks called by MACS2 at 54 hpf, ordered by fold-enrichment.

**Fig. S3.**
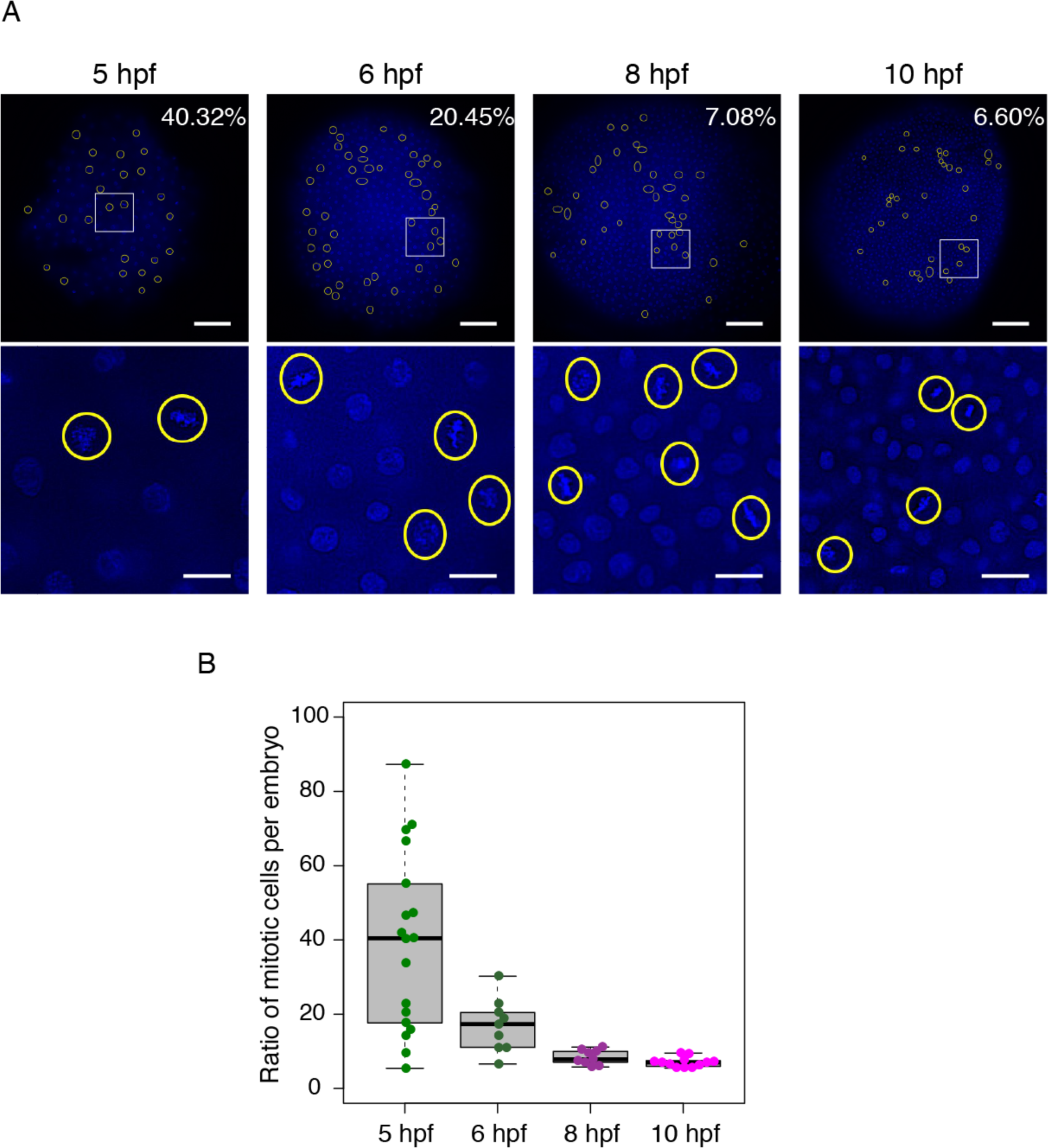
Cell cycle synchronization measured by counting mitotic cells per embryo. Anamniote embryos generally exhibit short and synchronous cell cycles until the blastula stage, followed by the transition into longer and asynchronous ones. In medaka, synchronous cell divisions take place with a cycle length of 30 to 35 minutes before the blastula stage. As mitotic chromosomes have unique 3D organization characterized by mega-base contact patterns, which may affect the global view of Hi-C contacts from whole embryos especially at early stages, we first evaluated the proportion of mitotic cells in morula (5 hpf) to pre-early gastrula (10 hpf) embryos by microscope observation. Collected embryos contained approximately 39%, 17%, 8%, and 7% of mitotic cells on average at 5 hpf, 6 hpf, 8 hpf and 10 hpf, respectively. We observed that cell division has already become asynchronous and the duration of interphase become longer at 5 hpf. Although the mixture of mitotic and interphase cells in an embryo prohibited us to remove mitotic chromatin from our analyses, we reasonably assumed that the majority of cells were at interphase cells, especially after 5 hpf. (A) Typical representations of counting mitotic cells before, during and after ZGA. The square windows at the top pictures were enlarged in the bottoms. The scale bars are 100 µm (top) and 20 µm (bottom). The yellow circles indicate mitotic cells. (B) Plots of the ratio of mitotic cells per embryo, showing that cell cycle synchronization already becomes less effective at 5 hpf.

**Fig. S4.**
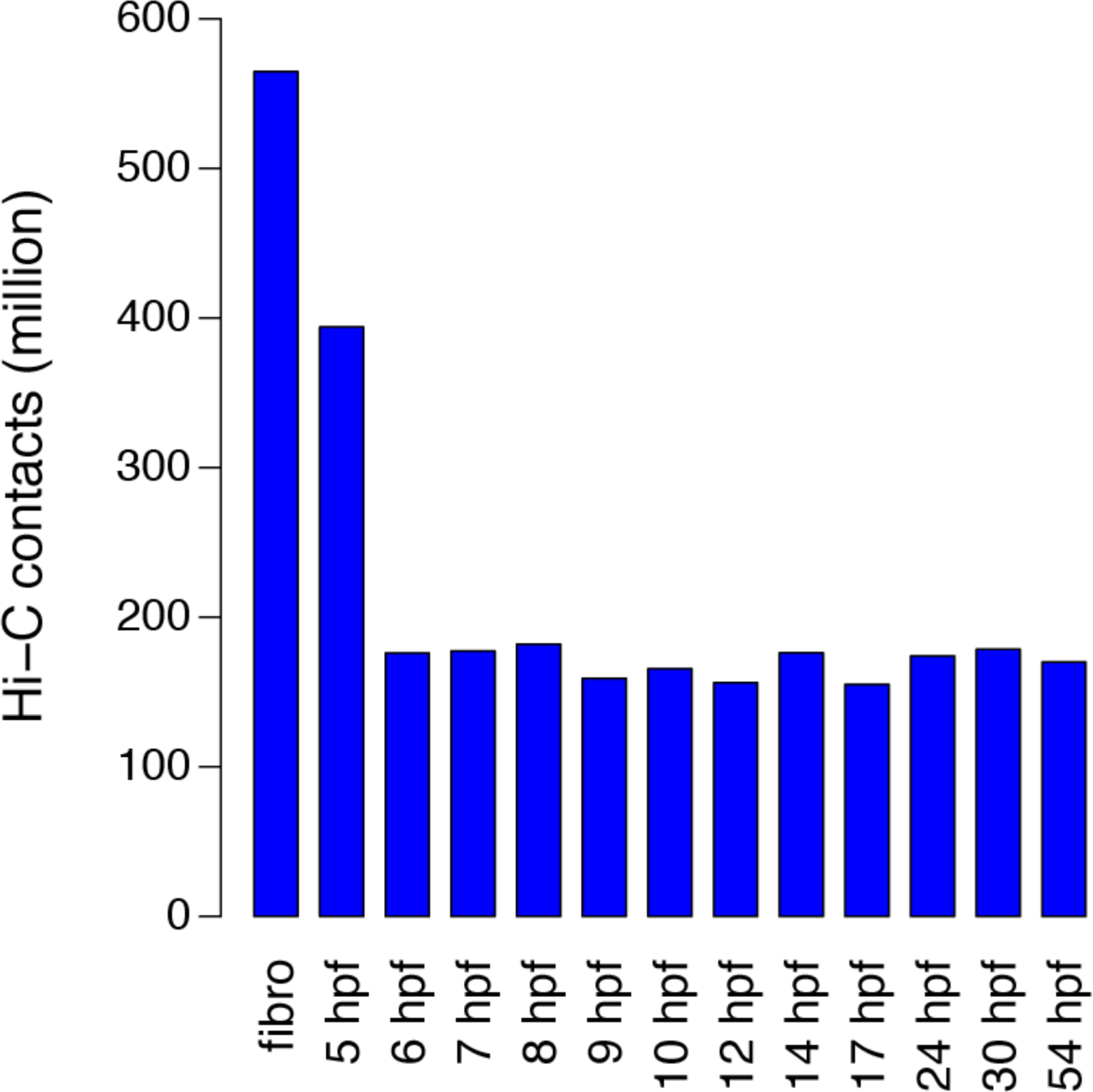
Number of Hi-C contacts across stages. The bargraph shows the number of Hi-C contacts at each stage after filtering using Juicer.

**Fig. S5.**
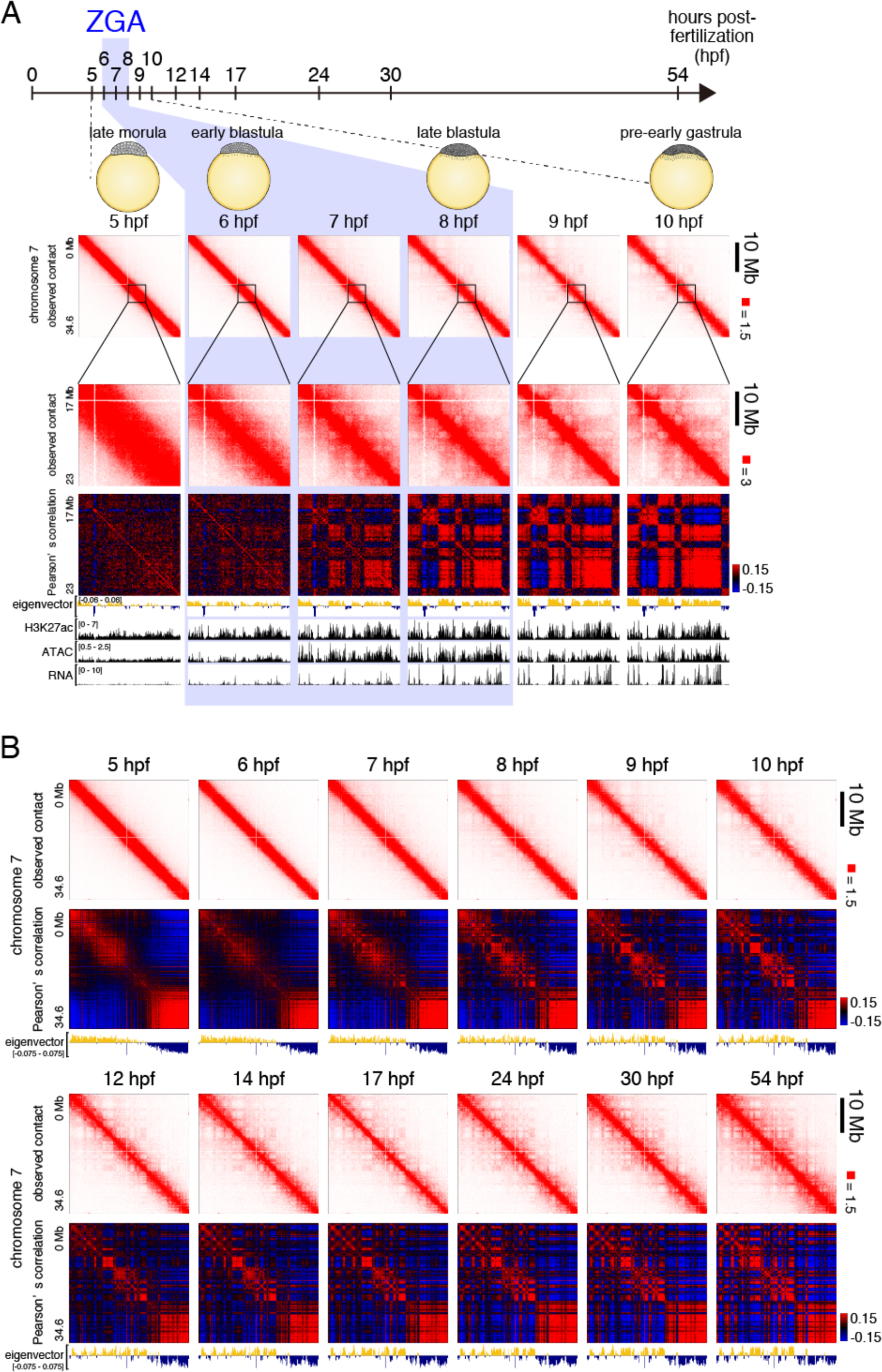
Compartmentalization through medaka development. (A) Compartmentalization at early developmental stages. Observed contact maps at 50 kb resolution are shown for whole chromosome 7 and enlarged views with corresponding Pearson’s correlation matrices, eigenvectors, H3K27ac ChIP-seq, ATAC-seq, and RNA-seq tracks for all indicated stages. For RNA-seq track, we assayed zygotic transcription. To facilitate the discrimination of maternal transcripts, which are ubiquitous in the egg throughout early development, and paternal transcripts, which were only derived from the paternal loci of the zygotic genome, we performed RNA-Seq on a cross between two medaka polymorphic inbred strains, d-rR and HNI. The rate of SNPs between the two strain is 2.5% (1 in 40 bp), making it possible to assign most reads to a paternal or maternal genome of origin. Using the level of paternal transcripts seen in RNA-seq as a proxy for overall zygotic transcription, we found that ZGA takes place between 6hpf and 8hpf. (B) Observed contact matrices, Pearson’s correlation matrices, eigenvector tracks are shown for whole chromosome 7 for all developmental stages used in this study.

**Fig. S6.**
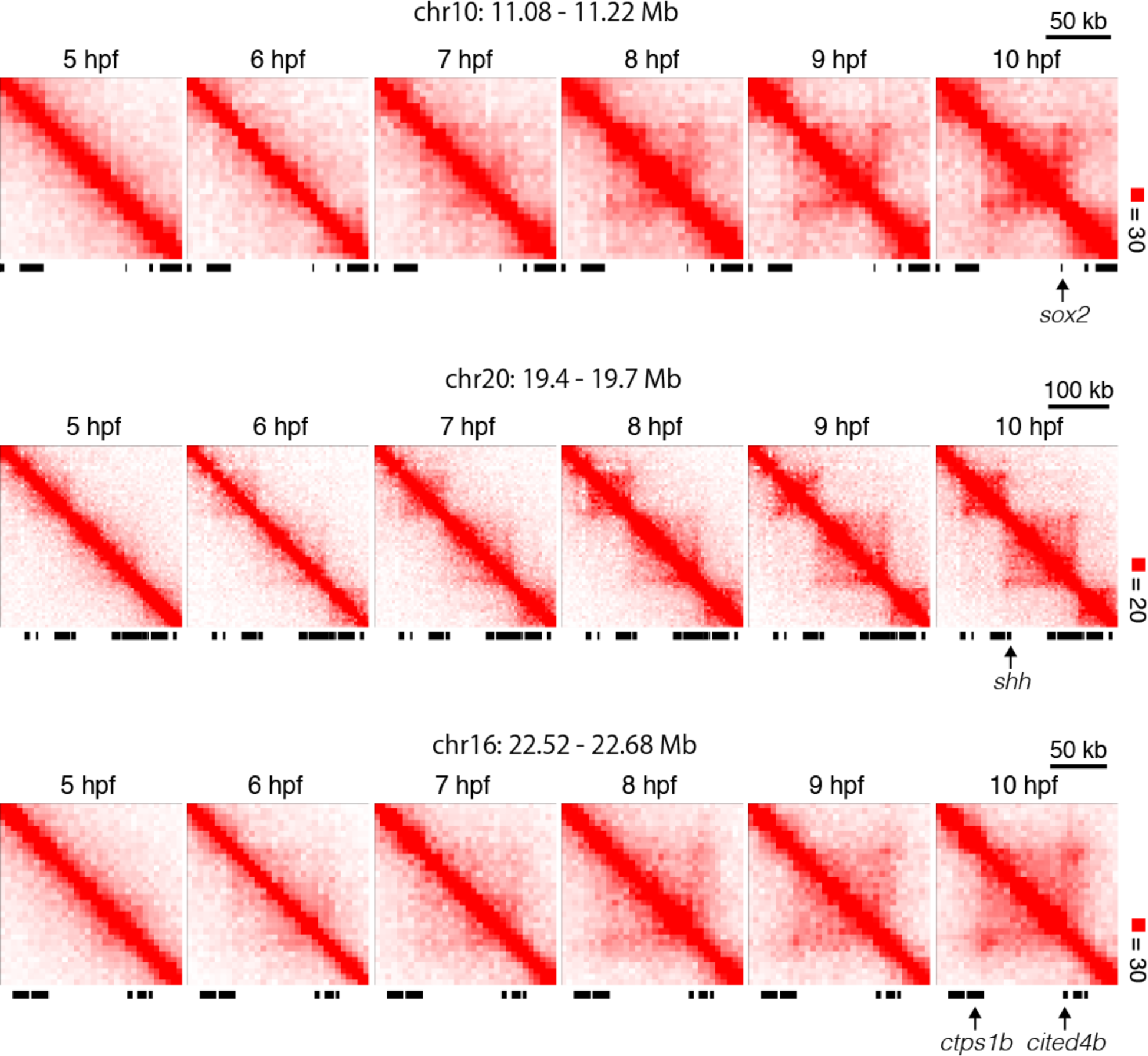
Examples of small contact domains emerging during ZGA. By manual inspection, we noticed some small contact domains which were overlooked by Arrowhead program because these contact domains were hard to detect within tight proximal contacts along the diagonals of contact matrices. Shown are several examples of relatively clear ones of those domains. These domains are absent at 5 hpf, but emerge during ZGA, and become clear at early gastrula stage (10 hpf).

**Fig. S7.**
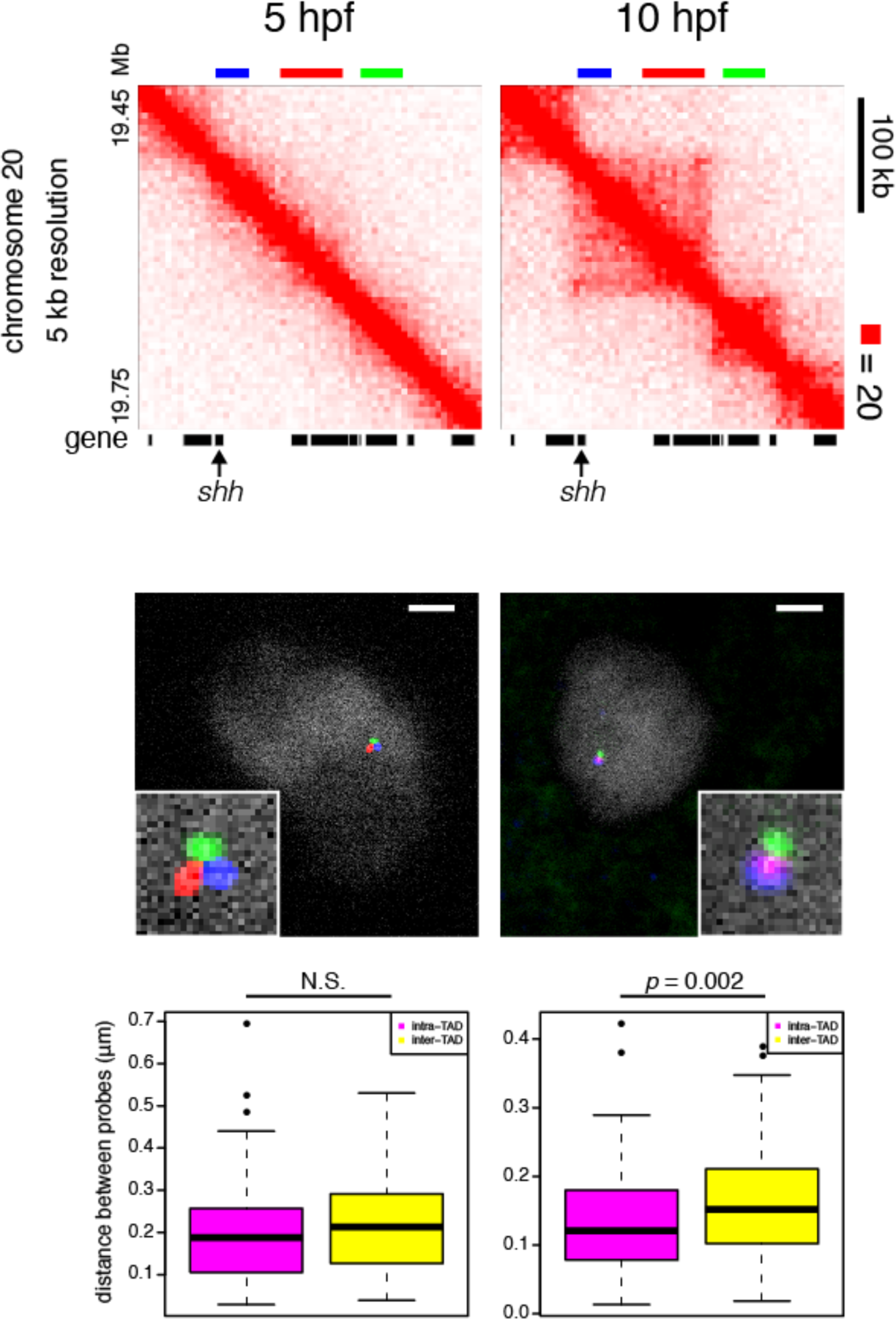
Confirmation of small contact domain at *shh* gene locus by microscopy. (top contact maps) A relatively small contact domain including *shh* gene was obviously established as development proceeded from 5 to 10 hpf (top). To validate the establishment of a small contact domain during ZGA *in vivo*, we performed 3D-FISH. Three probes were designed, two in a same contact domain to quantify an intra-TAD distance, and one in a neighboring contact domain to measure an inter-TAD distance. (middle microscope photos) We estimated the distance between a pair of two probes from a distance distribution among multiple microscope photos in interphase nuclei of 5 hpf and 10 hpf embryos. The representative views of their fluorescent signals were shown at the middle. The scale bar indicates 2 µm. (bottom box plots) Consistent with the Hi-C data, we observed no significant difference between the distance distributions between intra-TAD probes and inter-TAD probes at 5 hpf, but a significance increase in inter-TAD distance distribution at 10 hpf.

**Fig. S8.**
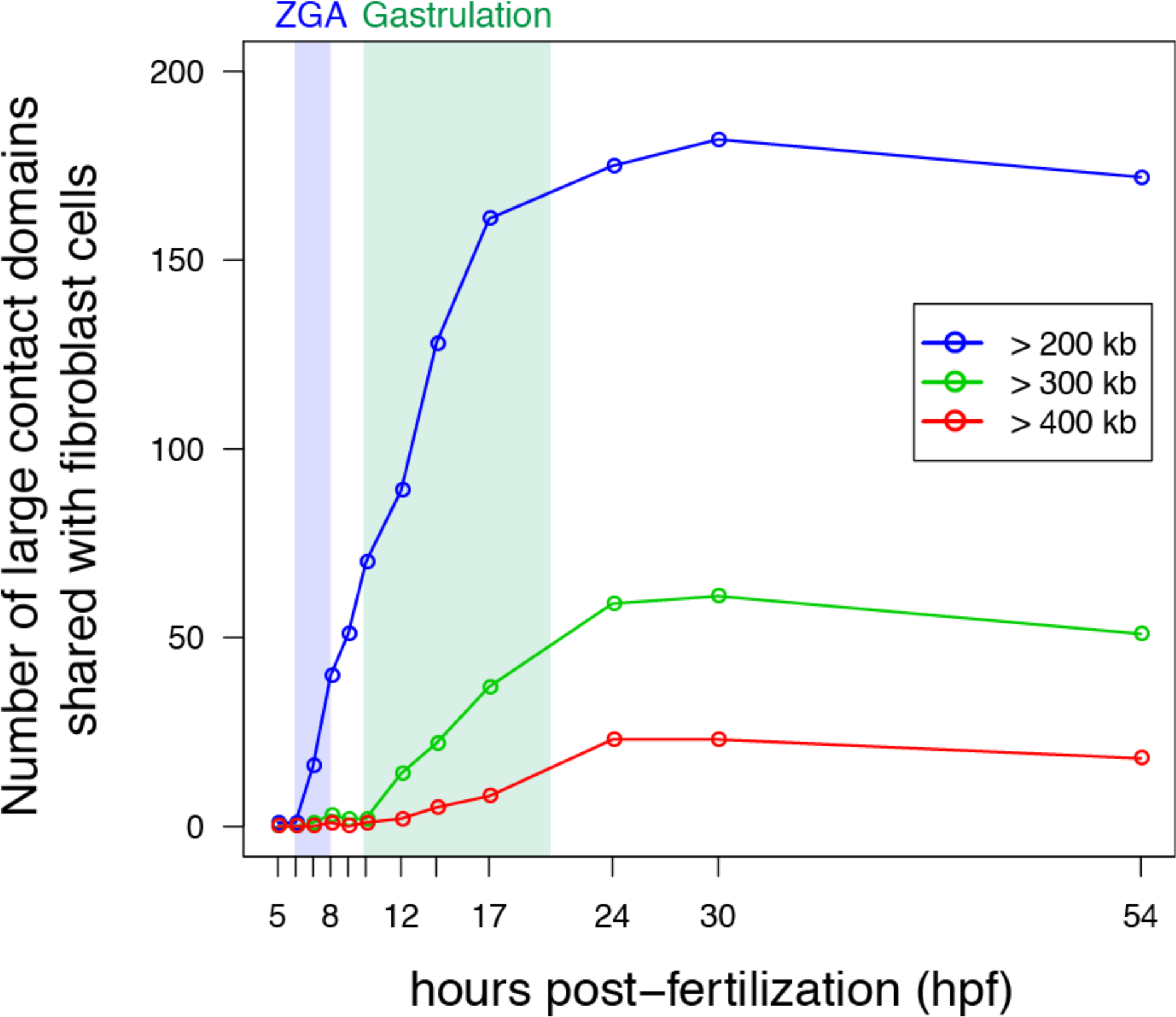
The majority of large contact domains associated with mature fibroblast cells emerge during gastrulation. The number of contact domains present in mature fibroblast cells is shown for different size ranges; ≥ 200 kb, ≥ 300 kb, and ≥ 400 kb.

**Fig. S9.**
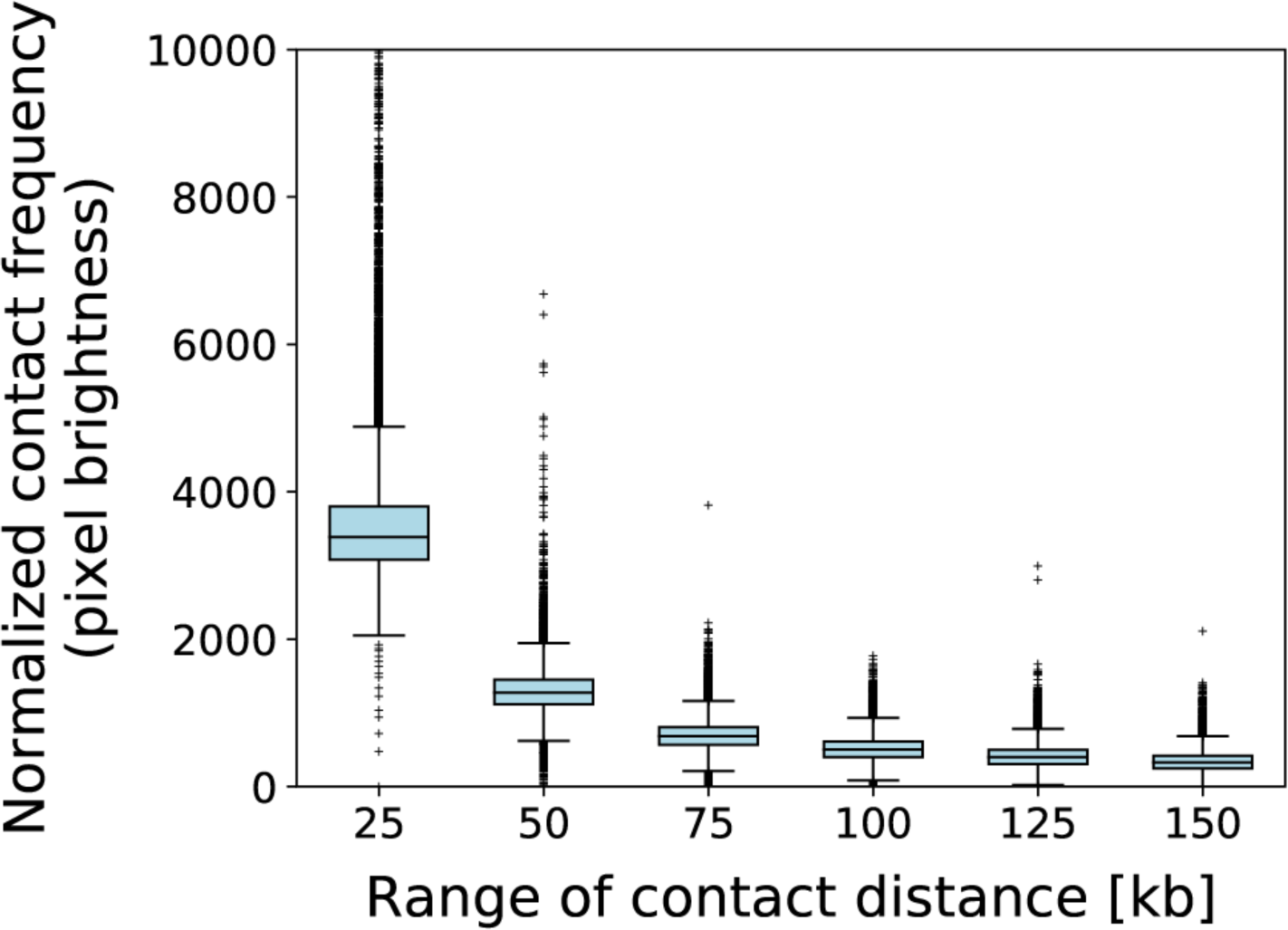
Frequency distribution of pixel brightness in the fibroblast contact frequency map. Each loop in a contact frequency map is identified as a local maximum peak in the region of size 35kb for 5kb resolution (50kb for 10kb resolution) from the center according to the peak calling method (9). It becomes harder to detect smaller loops that are closer to the diagonal of contact frequency map. To confirm this tendency, the box plots show the frequency distribution of pixel brightness in the fibroblast contact frequency map, where the brightness of a pixel represents a normalized contact frequency between pairs of positions at a distance. Distances are partitioned into ranges of length 25kb, which almost accords with the radius used for calling local maximum peaks. “25kb” in the x-axis, for example, shows the range smaller than 25kb, and “50kb” ranges from 25kb to 50kb. Of note, most of values in the 50kb box plot are smaller than the median in the 25kb box plot, and hence, many local maximum peaks in proximity to the diagonal are difficult to treat as loops. By contrast, highly bright pixels with extremely large contact frequencies in the 125kb box plot are also outliers in the 100kb box plot. Put another way, most of > 100kb loops are detectable as local maximum peaks, and hence, we focus on >100kb loops.

**Fig. S10.**
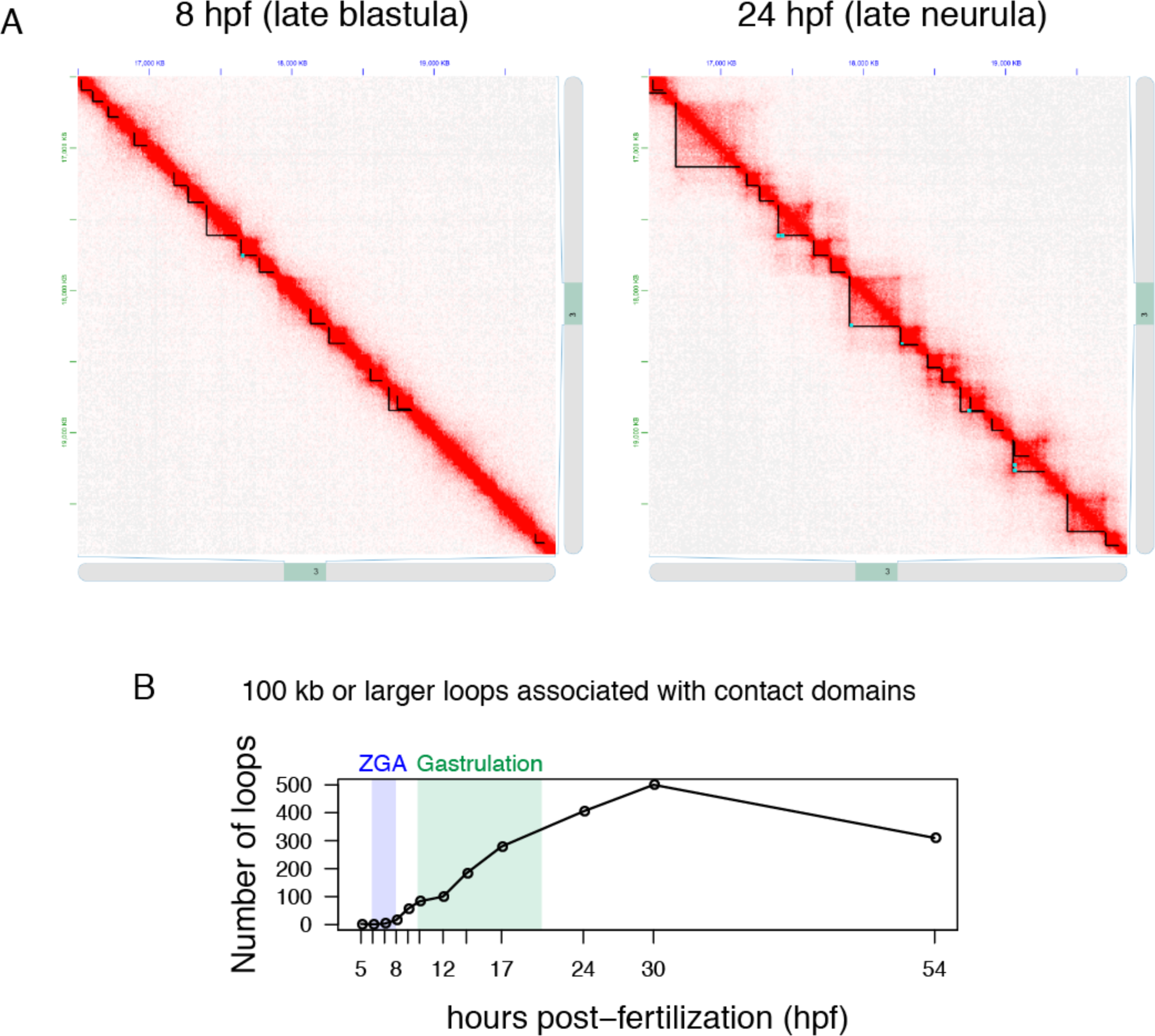
Number of loops detected across developmental stages. (A) Screen shots of Juicebox showing the representative loops and contact domains before gastrulation (8 hpf; late blastula) and after gastrulation (24 hpf; late neurula). (B) Loops smaller than 100 kb were difficult to distinguish from false positives due to the resolution limitation. Thus, we counted the number of loops (associated with contact domains) that are 100 kb or larger.

**Fig. S11.**
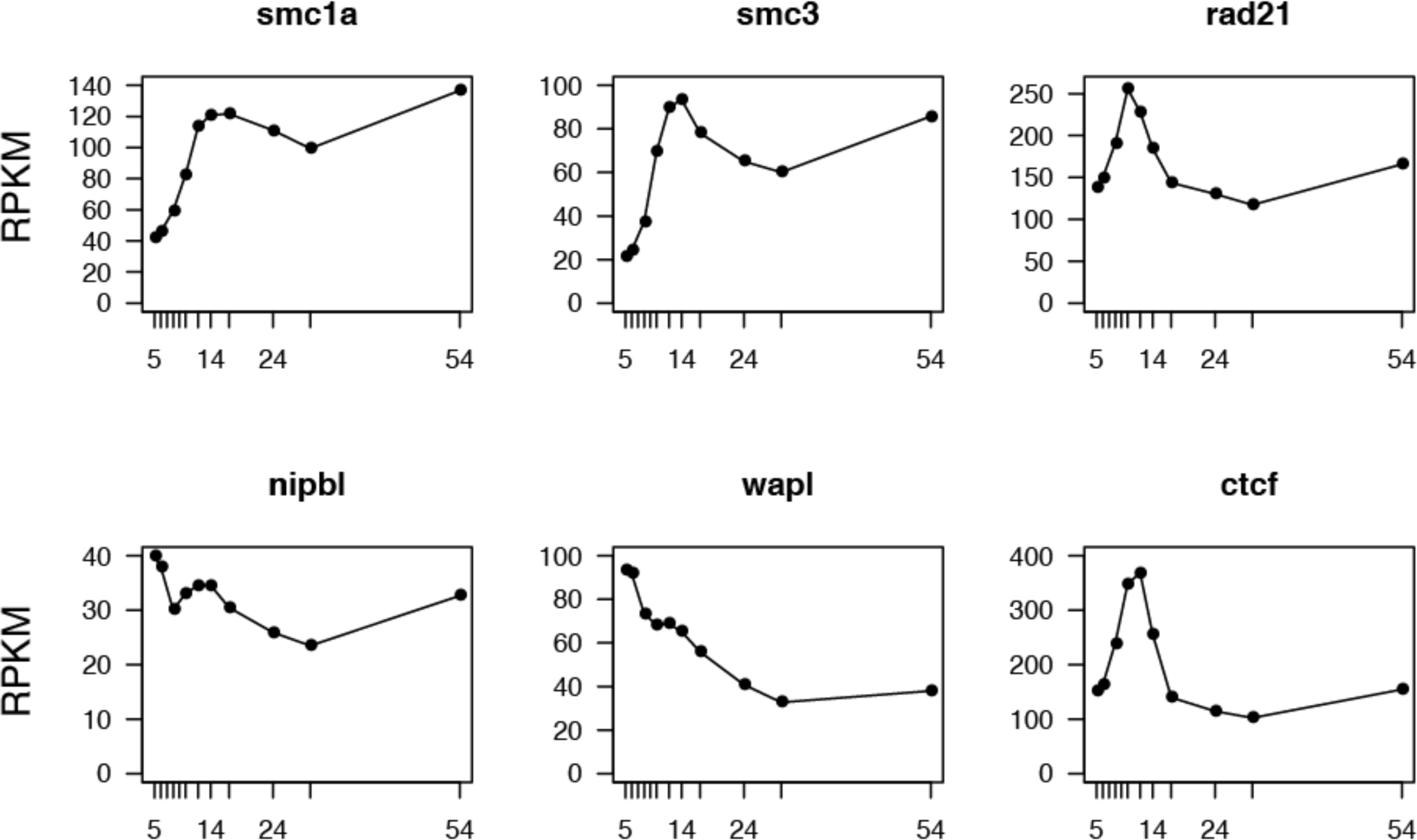
Abundance of transcripts associated with cohesin across development. The amounts of transcripts sampled from d-rR embryos were shown in RPKM through developmental stages. The genes were selected to be related to cohesin loop extrusion model; SMC1A, SMC3, and RAD21 proteins are cohesin subunits, NIPBL and WAPL proteins work for cohesion loading on and releasing from DNA respectively, and CTCF protein is a DNA binding protein which locates the loop domain boundaries co-localizing with cohesion complex.

